# The number of growing microtubules and nucleus-nucleus interactions uniquely regulate nuclear movement in *Drosophila* muscle

**DOI:** 10.1101/2020.04.22.054858

**Authors:** Mary Ann Collins, L. Alexis Coon, Riya Thomas, Torrey R. Mandigo, Elizabeth Wynn, Eric S. Folker

## Abstract

Nuclear movement is a fundamental process of eukaryotic cell biology. Skeletal muscle presents an intriguing model to study nuclear movement because its development requires the precise positioning of multiple nuclei within a single cytoplasm. Furthermore, there is a high correlation between aberrant nuclear positioning and poor muscle function. Although many genes that regulate nuclear movement have been identified, the mechanisms by which these genes act is not known. Using *Drosophila melanogaster* muscle development as a model system, and a combination of live-embryo microscopy and laser ablation of nuclei, we have found that phenotypically similar mutants are based in different molecular disruptions. Specifically, ensconsin (*Drosophila* MAP7) regulates the number of growing microtubules that are used to move nuclei whereas bocksbeutel (*Drosophila* emerin) and klarsicht (*Drosophila* KASH-protein regulate interactions between nuclei.

## INTRODUCTION

Since the identification of the Linker of Nucleoskeleton and Cytoskeleton (LINC) complex (Crisp et al., 2006; Starr and Fridolfsson, 2010; Tapley and Starr, 2013), the question of how nuclei move has been a pressing question in biology. The process of moving this heavy organelle is conserved throughout evolution in all cell types (Mosley-Bishop et al., 1999; Tran et al., 2001; Starr et al., 2001; Lee et al., 2002; Starr and Han, 2002; Del Bene et al., 2008; Zhang et al., 2009; Yu et al., 2011), thus magnifying the importance of understanding the underlying mechanism. Although many mechanisms have been described for mononucleated cells (Gundersen and Worman, 2013), how nuclei are moved in a syncytium has remained a mystery. Many genes that regulate nuclear position in syncytial skeletal muscle cells have been identified (Roman and Gomes, 2018), but how these genes contribute to nuclear movement and whether these genes regulate nuclear positioning through a single mechanism is not known.

In most contexts, nuclear movement is dependent on the microtubule cytoskeleton and its associated proteins which generate the force to move nuclei and the Linker of nucleoskeleton and cytoskeleton (LINC) complex which transmits force between the cytoskeleton and the nucleus. This is indeed true during the development of the syncytial abdominal musculature of *Drosophila melanogaster* embryos and larvae. Several microtubule associated genes including ensconsin/MAP7 (Metzger et al., 2012), Bsg25D/Ninein (Rosen et al., 2019), and the motors kinesin and cytoplasmic dynein (Folker et al., 2012, 2014) have been suggested to contribute to nuclear movement by regulating Kinesin activity (Metzger et al., 2012), microtubule stability (Rosen et al., 2019), and the application of force both directly on (Folker et al., 2014) and at a distance from (Folker et al., 2012). Similar experiments have shown that the LINC complex components klarsicht, (Elhanany-Tamir et al., 2012; Collins & Mandigo et al., 2017), Msp300 (Elhanany-Tamir et al., 2012), and klaroid (Tan et al., 2018) along with the emerin homologs bocksbeutel and Otefin (Collins & Mandigo et al., 2017; Mandigo et al., 2019) are also critical for nuclear positioning during muscle development. Despite identifying many of the factors that are critical for nuclear position, we know little about the mechanisms by which they support nuclear movement during muscle development.

The limited mechanistic understanding is in part driven by the complexity that many nuclei in a single cytoplasm creates. Although many studies investigating myonuclear movement have been done in cell culture (Cadot et al., 2012; Wilson and Holzbaur, 2012), such *in vitro* systems lack the complex signaling cascades that provide directionality cues to nuclei as they translocate, highlighting the importance of studying nuclear movement in an organismal context (Folker et al., 2014). Consequently, most *in vivo* work has relied on describing nuclei as mispositioned with little, if any, distinction between phenotypes (Metzger et al., 2012; Collins & Mandigo et al., 2017; Folker et al., 2012; Elhanany-Tamir et al., 2012). To better understand the mechanisms by which each gene regulates nuclear movement, it is critical to establish methods that can characterize nuclear phenotypes *in vivo* and distinguish between those that appear similar by a basic phenotypic scoring system. Here we describe a new analytical approach centered on live-embryo time-lapse microscopy and careful characterization of nuclear position combined with new tools to provide the first direct evidence that some factors necessary for nuclear movement are required to apply force to nuclei whereas other factors are necessary for the utilization of that force to reach a specific position rather than to move.

## RESULTS

### Disruption of *bocksbeutel* and *klarsicht* have distinct effects on myonuclear positioning compared to ensconsin in the *Drosophila* embryo

As a first approach, we have investigated the contributions of *bocksbeutel* (*Drosophila* emerin), *klarsicht* (*Drosophila* KASH-protein), and *ensconsin* (*Drosophila* MAP7). Each gene was zygotically removed in *Drosophila* embryos with the respective *bocks*^*DP01391*^ null (Collins & Mandigo et al., 2017), *klar*^*1*^ null (Welte et al., 1998), or *ens*^*swo*^ nonsense mutation (Metzger et al., 2012) alleles. Fixed images of *Drosophila* embryos showed that in controls, nuclei were in two clusters positioned at either end of the lateral transverse (LT) muscle whereas in *bocks*^*DP01391*^ and *klar*^*1*^ embryos, most of the nuclei were clustered together in a single group near the ventral end of the muscle (Fig. S1a), as we showed previously (Collins & Mandigo et al., 2017). Qualitatively, this clustering phenotype was similar to nuclear positioning defects observed in *ens*^*swo*^ embryos in which nuclei also failed to separate into distinct groups (Metzger et al., 2012). To quantitatively evaluate myonuclear position, the distance of each nuclear cluster from the dorsal and ventral muscle poles was measured. Since the LT muscles in all three mutants were significantly shorter (Fig. S2a, statistics summarized in Table S1), we measured the raw distance (Fig. S2) and the distance as percent of muscle length (Fig. S1). Compared to controls, nuclei in *bocks*^*DP01391*^ and *klar*^*1*^ embryos were positioned further from the dorsal muscle pole (Fig. S1b) yet closer to the ventral muscle pole (Fig. S1c), as previously described (Collins & Mandigo et al., 2017). However, nuclei in *ens*^*swo*^ embryos were positioned significantly further from both muscle poles when compared to controls or *bocks*^*DP01391*^ and *klar*^*1*^ embryos. Additionally, the distance between dorsal and ventral clusters was measured (Fig. S1d and Fig. S2d). The distance between clusters was significantly decreased in *bocks*^*DP01391*^ and *klar*^*1*^ embryos because distinct clusters of nuclei formed in only a small fraction of muscles (Fig. S2e and f). In contrast, since nuclei failed to separate in nearly all *ens*^*swo*^ muscles, this distance was approximately 0 µm. Finally, we measured the area of dorsal and ventral clusters to compare the distribution of nuclei as previously described (Collins & Mandigo et al., 2017). In controls, nuclei were evenly distributed between the two clusters, whereas more nuclei remained associated within the ventral cluster in *bocks*^*DP01391*^ and *klar*^*1*^ embryos, thus significantly decreasing the nuclear separation ratio (Fig. S1e), consistent with previous data (Collins & Mandigo et al., 2017). Similarly, in the rare case in which nuclei separated in *ens*^*swo*^ embryos, there were more nuclei in the ventral cluster compared to the dorsal cluster. Although the total area occupied by nuclei was similar between controls, *bocks*^*DP01391*^, and *klar*^*1*^, it was significantly reduced in *ens*^*swo*^ embryos (Fig. S2i). However, the number of nuclei was the same between controls and *ens*^*swo*^ embryos, indicating that fusion is not affected (Fig. S3a and b). Additionally, the total volume occupied by nuclei is the same in both genotypes (Fig. S3c and Movies S1 and S2). Thus, the reduced area is due to nuclei occupying a greater depth in the *ens*^*swo*^ embryos.

Based on these measurements, the most dominant phenotype observed in control embryos was nuclei that separated into two distinct groups of equal size. In *bocks*^*DP01391*^ and *klar*^*1*^ embryos, nuclei either remained as a single cluster positioned near the ventral end of the muscle (Fig. S1f and g, “clustered” and “spread”) or in two clusters in which the dorsal group was significantly smaller than the ventral group (Fig. S1f and g, “separated: unequal distribution”). Finally, the most dominant phenotype observed in *ens*^*swo*^ embryos was a single cluster positioned near the center of the muscle (Fig. S1f and h, “swoosh”). In total, these data indicate that while bocksbeutel, klarsicht, and ensconsin are all required for proper nuclear movement, the disruption of *ens* causes a distinct type of nuclear positioning defect compared to the disruption of *bocks* and *klar* and suggest that these genes may regulate distinct aspects of nuclear movement.

### *Ensconsin* is necessary for nuclear movement whereas *bocksbeutel* and *klarsicht* are necessary to separate nuclei

To investigate these phenotypes further, the position of nuclear clusters within the LT muscles was tracked over the course of 2 hours. In control muscles, once all nuclei separated into two distinct clusters, these clusters migrated toward opposite muscle ends, steadily increasing the distance between themselves (Fig. 1a and Movie S3, left panel). However, 100% of all nuclei observed in *ens*^*swo*^ muscles failed to separate over the time course (Fig. 1a, yellow brackets and Movie S6, left panel), significantly reducing the separation speed to 0 μm/hr (Fig. 1b and c). Similarly, nuclei that remained associated together in *bocks*^*DP01391*^ and *klar*^*1*^ muscles also failed to separate (Fig. 1b blue data points and Movies S4 and S5, left panels). However, this non-separation phenotype was only observed in approximately 50% of muscles (Fig. 1c). In the other 50% of muscles, a single nucleus separated and migrated towards the dorsal end of the muscle (Fig. 1a, yellow arrows), at a rate slightly faster than control nuclei (Fig. 1b, gray data points). Furthermore, the morphology of the single clusters was different in *bocks*^*DP01391*^ and *klar*^*1*^ compared to *ens*^*swo*^. In *ens*^*swo*^ clustered nuclei were spherical, whereas nuclear clusters in *bocks*^*DP01391*^ and *klar*^*1*^ embryos were significantly elongated (Fig. 1d).

**Fig. 1.**
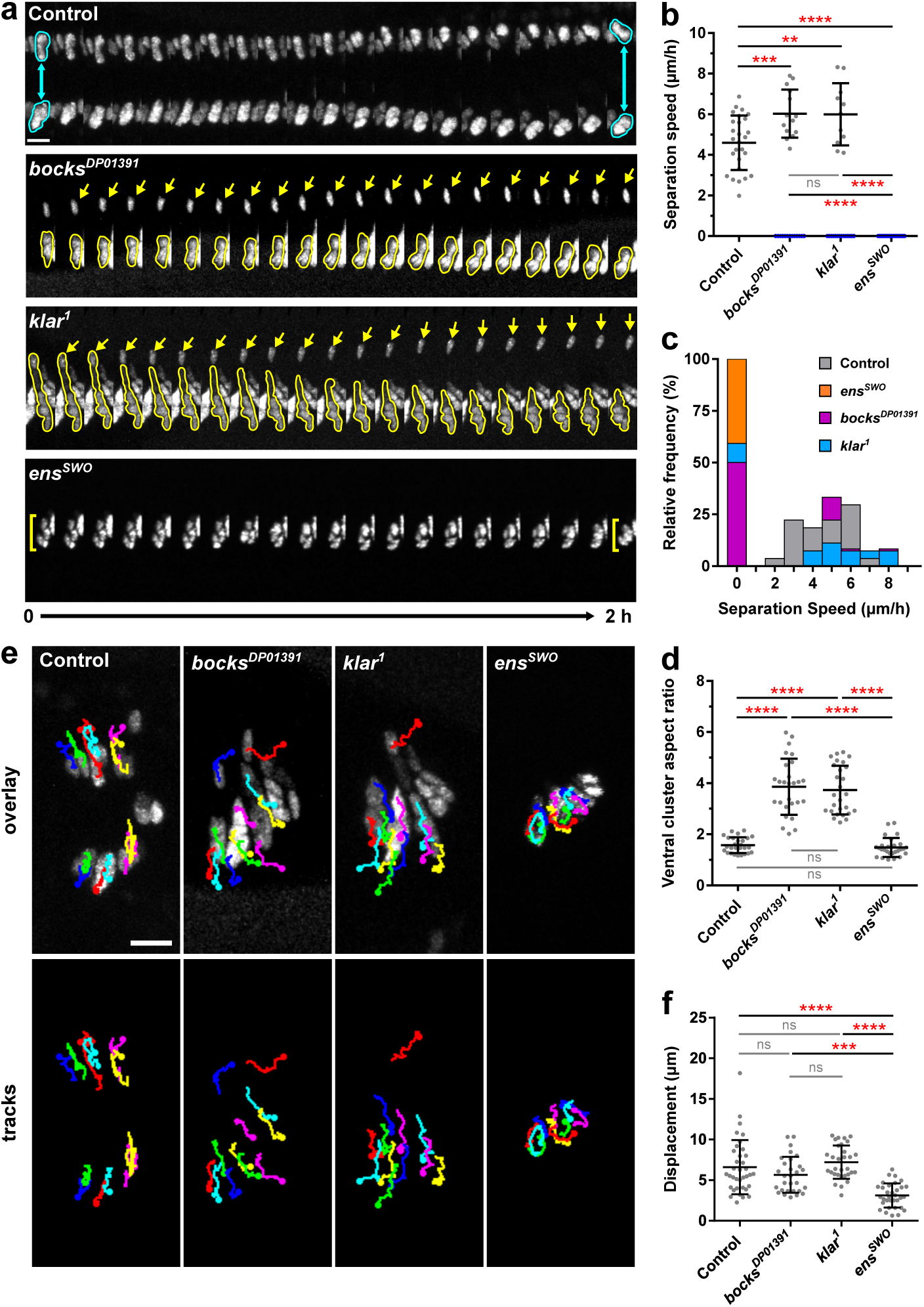
Bocksbeutel, klarsicht, and ensconsin are necessary for the proper separation of myonuclei in *Drosophila* embryos. (**a**) Montages from time-lapse acquisitions showing the separation of the dorsal cluster from the ventral cluster of nuclei within a single lateral transverse (LT) muscle of a stage 15 (15 hours AEL) embryo for the indicated genotypes. Nuclei outlined in cyan indicate the proper separation of nuclei into two distinct clusters (control). Yellow arrows indicate an escaper nucleus that separates from the ventral group in either *bocks*^*DP01391*^ or *klar*^*1*^ mutant embryos. Yellow brackets indicate nuclei that fail to separate and remain associated as a single cluster (*ens*^*swo*^). Scale bar, 10 µm. (**b**) The separation speed of nuclear clusters. Data points correspond to the speed measured from a single LT muscle. Gray data points indicate the speed at which the dorsal and ventral clusters of nuclei separate from one another, whereas blue data points indicate nuclei that failed to separate (speed = 0 µm/h). Error bars indicate the s.d. from ≥25 LT muscles for each genotype taken from independent experiments. (**c**) The relative distribution of nuclear separation speeds. (**d**) The aspect ratio of the ventral nuclear cluster measured at 0 h. Data points correspond to the ventral nuclear cluster within a single LT muscle. Error bars indicate the s.d. from ≥25 LT muscles for each genotype taken from independent experiments. (**e**) Tracks following the movement of individual nuclei within four LT muscles over the course of two hours, superimposed over the first frame (t = 0 h). Scale bar, 10 µm. (**f**) The displacement of individual nuclei. Data points correspond to the displacement of a single nucleus. Error bars indicate the s.d. from 36 nuclei for each genotype taken from three independent experiments. For (b), (d), and (f), One-way ANOVA with Tukey HSD post hoc test was used to assess the statistical significance of differences in measurements between all experimental groups.

The trajectory of individual myonuclei within each cluster was then tracked over the 2-hour time course (Fig. 1e). The total displacement of nuclei in *bocks*^*DP01391*^ and *klar*^*1*^ embryos was similar to controls, even in ventral cluster where more nuclei were present (Fig. 1f and Movies S3-5, right panels). Although the displacement was similar to controls, all of the nuclei within the cluster moved ventrally. However, nuclei that did stochastically separate from the ventral cluster did migrate dorsally suggesting that the interactions between nuclei within a cluster is restricting the movement toward the ventral end of the muscle. Conversely, the displacement of nuclei in *ens*^*swo*^ embryos was significantly decreased, as nuclei rotated within in the cluster but did not translocate (Fig. 1f and Movies S6, right panel). Together these data suggest that in *ens*^*swo*^ mutants, the ability of the cell to exert force on nuclei is reduced. However, the movement and subsequent displacement of the nuclei in the *klar*^*1*^ and *bocks*^*DP01391*^ suggests that force production is normal and that instead nuclei are being actively maintained in a single cluster.

### Laser ablation of myonuclei demonstrates that the application of force onto nuclei is *ensconsin*-dependent

The fact that the nuclei were elongated in *bocks*^*DP01391*^ and *klar*^*1*^ mutants compared to controls suggested that they may be under tension. To test this hypothesis, we used 2-photon laser ablation to remove individual nuclei and measure the response of the neighboring nuclei within the syncytium (Fig. S4a). When a nucleus was ablated in controls (1 s, yellow circle and Movie S7), the remaining nuclei within the cluster moved away from the ablation site, toward the center of the muscle fiber (Fig. S4d, 2–5 s). Nuclei in the opposite cluster also moved towards the muscle center. However, the nuclei in the neighboring LT muscles did not respond to the ablation. Furthermore, ablation did not affect the health of the muscle or the animal. Imaging of the transmitted light demonstrated that there was no gross damage to the embryo. Furthermore, three hours after ablation, nuclei returned to their proper position adjacent to the muscle end. Similar movements of nuclei have been seen to occur due to muscle contractions, but in these cases, muscles detached from the tendon and formed a spheroid from which neither the muscle morphology or the nuclear position recovered (Auld et al., 2018b). Thus, the return of nuclei to the end of the muscle is consistent with the nuclei moving and not movement of the muscle ends due to contractions. Finally, ablation did not affect viability as embryos were able to developmentally progress to stage 17, initiate muscle contraction and hatching (Fig. S4e), and crawl out of the field of view.

We then ablated nuclei in muscles of animals where nuclei had failed to separate into distinct clusters (Fig. 2a). When compared to controls, the area of the ventral clusters in *bocks*^*DP01391*^ (Movie S8) and *klar*^*1*^ (Movie S9) embryos before ablation was significantly larger (Fig. 2b, before). After ablation, the remaining nuclei moved away from the ablation site and showed a 43% reduction in size in both genotypes (Fig. 2b, b’ after). The dramatic decrease in size suggests that the stretching of nuclei, in addition to the greater number of nuclei present, contributed to the difference in the size of the clusters. In contrast, nuclei in *ens*^*swo*^ embryos (Movie S10) moved only slightly after ablation (Fig. 2a) and their size was reduced by only 10%, a value consistent with the removal of 1 out of 6-7 nuclei (Fig. 2b, b’). In addition, after ablation, clusters in *bocks*^*DP01391*^ and *klar*^*1*^ embryos traveled a greater distance compared to controls while clusters in *ens*^*swo*^ embryos traveled a shorter distance (Fig. 2c and c’). Similarly, the clusters in *bocks*^*DP01391*^ and *klar*^*1*^ had a greater initial velocity compared to controls whereas nuclei in *ens*^*swo*^ embryos had a reduced initial velocity (Fig. 2d and d’). Together, these data demonstrate that nuclei in *bocks*^*DP01391*^ and *klar*^*1*^ embryos are under more tension than nuclei in controls, while nuclei in *ens*^*swo*^ embryos are under less tension. This is consistent with the hypothesis that ensconsin is necessary for the application of force to nuclei but that klarsicht and bocksbeutel are necessary for the directed movement of nuclei in response to that force.

**Fig. 2.**
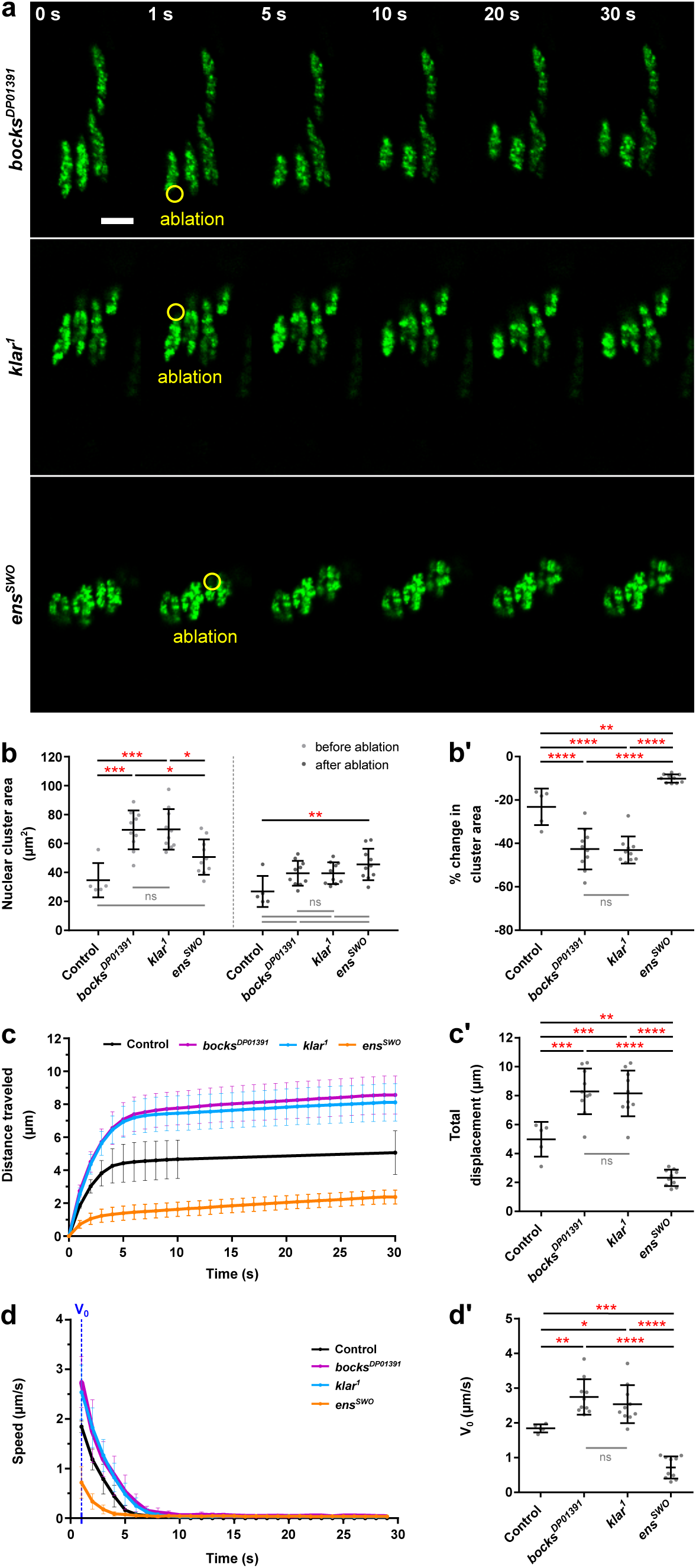
Nuclei in *bocksbeutel* and *klarsicht* mutants are under more tension than nuclei in *ensconsin* mutants. (**a**) Montages from time-lapse acquisitions showing the ablation of a myonucleus within the lateral transverse muscles of a stage 16 (16 hours AEL) embryo for the indicated genotypes. The first frame shows the nuclei before ablation (0 s). The next frame (1 s) shows the ablation of a single nucleus (yellow circle), followed by the subsequent response of the remaining nuclei after ablation (5-30 s). Scale bar, 10 µm. (**b**) The average area of nuclear clusters before and after ablation. (**b’**) The same data in (b) represented as a percent change in cluster area. A negative change in area indicates that the size of the nuclear cluster decreased after the ablation. (**c**) The average displacement of nuclear clusters after ablation as a function of time. (**c’**) The average total displacement of nuclear clusters after ablation. (**d**) The average change in speed of nuclear clusters after ablation as a function of time. (**d’**) The average initial speed (V_0_) of nuclear clusters the first second after ablation. Data points in (b–d’) correspond to an individual ablation event. Error bars indicate the s.d. from ≥5 ablation events performed in different embryos for each genotype. One-way ANOVA with Tukey HSD post hoc test was used to assess the statistical significance of differences in measurements between all experimental genotypes to controls.

### Loss of *bocksbeutel* and *klarsicht*, and ensconsin are required for the organization of microtubules in *Drosophila* larval skeletal muscle

Since myonuclei are physically linked to the microtubule cytoskeleton (Tassin et al., 1985; Espigat-Georger et al., 2016), ensconsin is a microtubule binding protein (Bulinski and Bossler, 1994; Gallaud et al., 2014), and nuclear envelope proteins have been demonstrated to impact microtubule organization (Hale et al., 2008; Bugnard et al., 2005; Starr and Fridolfsson, 2010; Gimpel et al., 2017), we hypothesized that the differences in nuclear behaviors may be linked to variations in microtubule organization. For this analysis we used larvae in which the muscles are 100X larger and therefore provide greater resolution of microtubule organization. Additionally, we used the ventral longitudinal muscle 3 (VL3) of stage L3 larvae (Fig. 3a), which are a large, flat, rectangular muscle group that is at the top of a dissected larva. We focused on two distinct regions of microtubules that are uniquely organized. The first region pertained to areas of the muscle, distant from nuclei, where microtubules intersect to form a lattice (Fig. 3a, yellow box, and 3b) while the second region was adjacent to nuclei and consisted of microtubules that emanate directly from the nuclei (Fig. 3a, cyan box, and 3c). As previously reported (Collins & Mandigo et al., 2017; Elhanany-Tamir et al., 2012), nuclei in *bocks*^*DP01391*^ and *klar*^*1*^ larvae were mispositioned in a single row along the anterior-posterior axis of the muscle compared to nuclei in controls, which were evenly distributed in two parallel lines. Analysis of the lattice network of microtubules (Fig. 3b) was performed using the Texture Detection Technique (TeDT), which detects the angles at which neighboring microtubules intersect (Liu and Ralston, 2014). In controls, the dominant intersection angles were parallel (0°, 180°, 360°) to the anterior-posterior axis of the muscle (Fig. 3d, average in Fig. 3d’). Microtubules in *bocks*^*DP01391*^, *klar*^*1*^, and *ens*^*swo*^ larval muscles were highly disorganized, with an overall reduction in the frequency of microtubules intersecting at every 180° (Fig. 3d’).

**Fig. 3.**
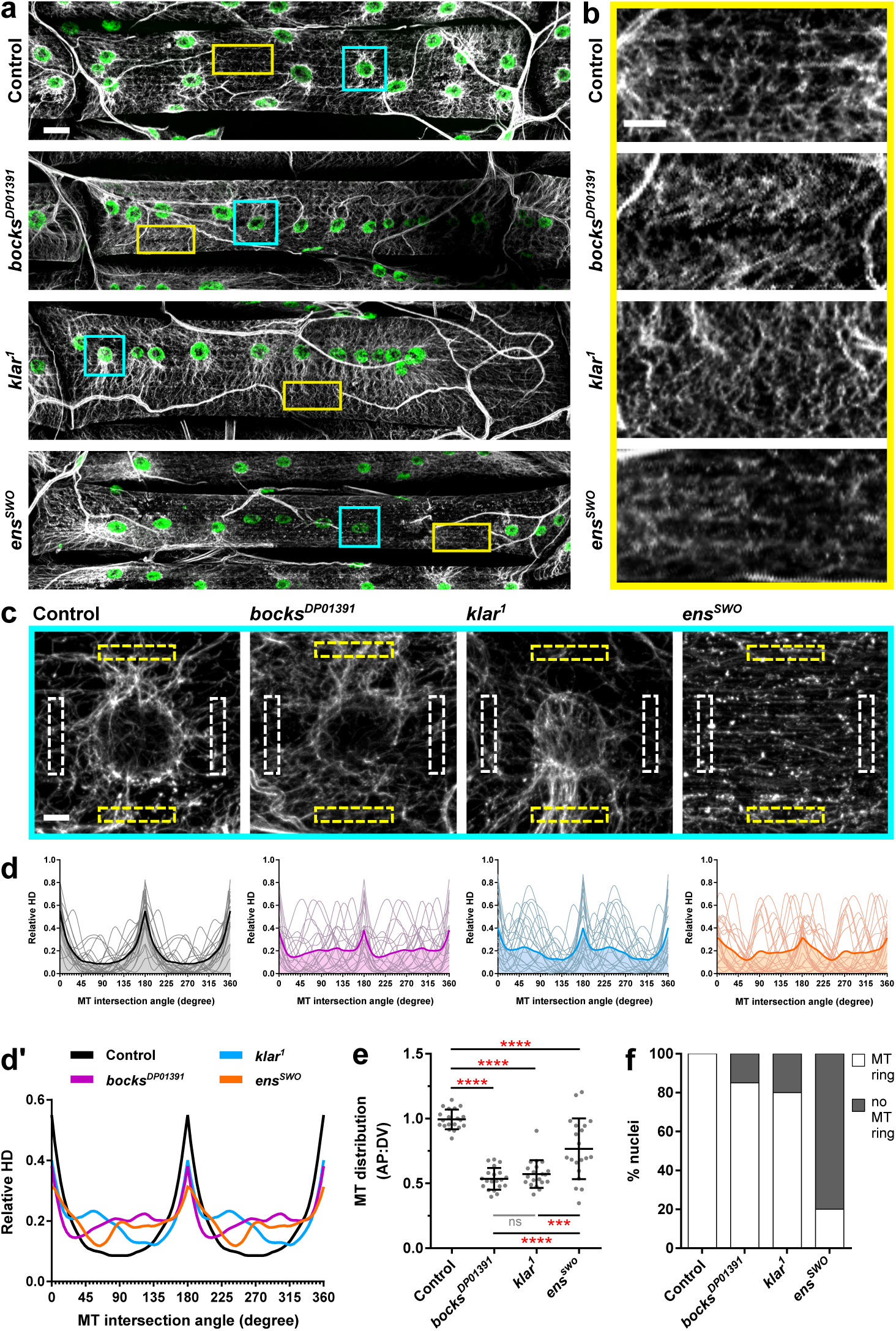
*Bocksbeutel, klarsicht*, and *ensconsin* disrupt microtubule organization in *Drosophila* larval skeletal muscle. (**a**) Immunofluorescence images of ventral longitudinal muscle 3 from stage L3 larvae for the indicated genotypes. Microtubules (α-tubulin) in gray, myonuclei in green. Scale bar, 25 µm. (**b**) Magnified regions of the microtubule lattice taken from the images shown in (a), as indicated by the yellow box. Scale bar, 10 µm. (**c**) Magnified regions of microtubules emanating from myonuclei taken from the images shown in (a), as indicated by the cyan box. White dotted boxes indicate the location of anterior and posterior fluorescence intensity measurements while yellow dotted boxes indicate the location of dorsal and ventral fluorescence intensity measurements for microtubule polarity analysis. Scale bar, 5 µm. (**d**) TeDT analysis of microtubule lattice regions. Intersection angles are represented as directional histograms (HD) from 0° to 360°. Thin lines indicate TeDT analysis for individual MT lattice regions, while the thick color line indicates the average of 20 MT lattice regions for each genotype. (**d’**) The average TeDT analysis from 20 MT lattice regions as shown in (d) for *bocks*^*DP01391*^ (purple), *klar*^*1*^ (blue), and *ens*^*swo*^ (orange) compared to controls (black). (**e**) The polarity of microtubules around myonuclei, represented as the microtubule distribution ratio for each nucleus. Data points correspond to the microtubule distribution ratio of a single nucleus. Error bars indicate the s.d. from 20 nuclei for each genotype from ≥10 VL3 muscles. One-way ANOVA with Tukey HSD post hoc test was used to assess the statistical significance of differences in measurements between all experimental groups. (**f**) The frequency in which microtubule rings were observed around nuclei in each of the indicated genotypes. A total of 20 nuclei were analyzed for each genotype from ≥10 VL3 muscles.

To evaluate the organization of microtubules that extend off of nuclei, we counted the percentage of nuclei that have a dense ring of microtubules on the nuclear periphery (Fig. 3f) and measured the proportion of microtubules on the dorsal-ventral axis of the muscle versus the anterior-posterior axis (Fig. 3e). In controls, all nuclei had a ring of microtubules and the distribution ratio was close to 1.0, indicating that microtubules are uniformly emanating from nuclei. Although 85% of *bocks*^*DP01391*^ and 80% of *klar*^*1*^ nuclei had a ring of microtubules (Fig. 3f), the distribution ratio was reduced to 0.535 and 0.572 in *bocks*^*DP01391*^ and *klar*^*1*^ larvae respectively (Fig. 3e), indicating that more microtubules are extending along the dorsal-ventral axis compared to the anterior-posterior axis. However, only 20% of nuclei in *ens*^*swo*^ mutants had rings (Fig. 3f) and there was a wide distribution in the proportion of microtubules on the dorsal-ventral and anterior-posterior axes compared to both controls, *bocks*^*DP01391*^, and *klar*^*1*^ mutants (Fig. 3e). Together, these data indicate that although bocksbeutel, klarsicht, and ensconsin are necessary to maintain the link between myonuclei and microtubules, the disruption of *bocks* or *klar* results in the reorganization of microtubules around mispositioned nuclei whereas the disruption of *ens* completely disrupts the general organization of microtubules throughout the muscle.

Our finding that microtubule organization is dependent on ensconsin differs from previous studies that suggested that the function of ensconsin was only to activate Kinesin (Barlan et al., 2013). To determine whether the disruption in microtubule organization was a consequence of mispositioned nuclei or a contributor to nuclear movement, we examined the behavior of EB1 during embryonic muscle development when nuclei are actively moving. EB1 comets were tracked for 1 minute in the LT muscles (Movies S11 and S12) and the dorsal oblique (DO) muscles (Movies S13 and S14), a set of broad, flat muscles that are more amenable to fast, live-embryo imaging (Fig. 4a). The location from which EB1 emerged, their direction of travel, and their speed was the same in controls and *ens*^*swo*^ embryos in both muscle types (Fig. 4b and c). However, the number of EB1 comets was significantly decreased in both LT and DO muscles of *ens*^*swo*^ embryos (Fig. 4e) indicating that ensconsin is critical to regulating the number of growing microtubules during *Drosophila* muscle development. Because most microtubules emanate from the nuclei in *Drosophila* larval muscles, the decrease in microtubule number (Fig. 4e) is consistent with the decreased percentage of nuclei with microtubule rings (Fig. 3f), further supporting a role for ensconsin in maintaining the general organization of microtubules within skeletal muscles.

**Fig. 4.**
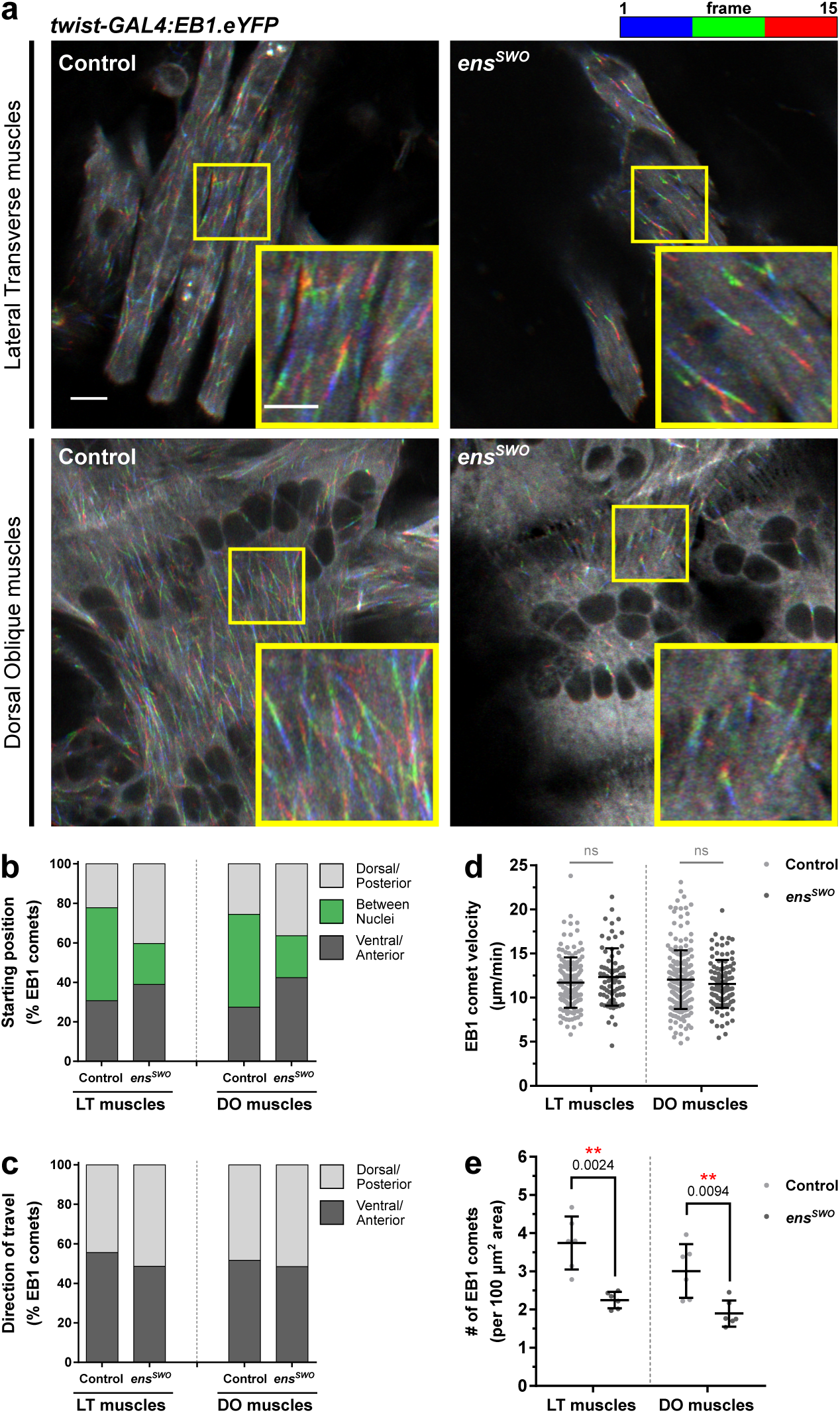
Depletion of *ensconsin* decreases the number of EB1 comets in *Drosophila* embryonic muscles. (**a**) Temporal overlays tracking EB1 comets for 15 s in the lateral transverse (LT) muscles and dorsal oblique (DO) muscles of stage 16 control and *ens*^*swo*^ embryos. Scale bar, 5 μm. (inset in yellow box) Magnified regions of the temporal overlays tracking EB1 comets for 15 s. Scale bar, 3 μm. (**b**) The frequency of EB1 comets observed in controls and *ens*^*swo*^ muscles starting in the dorsal/posterior muscle pole region, ventral/anterior muscle pole region, or the region between nuclei. (**c**) The frequency of EB1 comets observed in controls and *ens*^*swo*^ muscles traveling either toward the dorsal/posterior muscle pole or the ventral/anterior muscle pole. (**d**) The average velocity of EB1 comets in controls and *ens*^*swo*^ muscles. Data points correspond to the velocity measured from a single EB1 comet. Error bars indicate the s.d. from EB1 comets measured from 6 different embryos for each muscle group taken from independent experiments. (**e**) The average number of EB1 comets counted in controls and *ens*^*swo*^ muscles, normalized to the muscle area. Data points correspond to the total number of EB1 comets counted from a single embryo. Error bars indicate the s.d. from 6 different embryos for each muscle group taken from independent experiments. For (d) and (e), Student’s t-test with Welsh’s correction was used to assess the statistical significance of differences in measurements between ensconsin-depleted embryos and controls for each muscle group.

## DISCUSSION

All together, these data demonstrate that nuclear movement in a muscle syncytium requires both the transmission of force from the cytoskeleton to the nucleus and the separation of nuclei from their neighbors (Fig. 5). Disruption of these two separate processes produces superficially similar nuclear positioning phenotypes, but careful analysis of the precise position, shape, and movement of nuclei clearly indicates that there are distinct molecular underpinnings. Consistent with this, we found that loss of ensconsin contributes to the application of force to nuclei by regulating the number of growing microtubules. Surprisingly, force was applied to nuclei in the absence of the KASH-domain protein klarsicht or the emerin homolog bocksbeutel. Consequently, nuclei moved a similar total distance to those nuclei in control embryos. However, nuclei remained attached rather than separating and therefore were all moved toward the ventral end of the muscle. Interestingly in *bocks*^*DP01391*^ and *klar*^*1*^ mutants, nuclei did rarely separate from the single cluster and move as individuals to the dorsal end of the muscle. This observation is consistent with the phenotype being based in aberrant associations between nuclei and not a disruption of directional cues. Finally, we use laser ablation of individual nuclei to demonstrate that nuclei in *bocks*^*DP01391*^ and *klar*^*1*^ mutants are under increased tension compared to controls whereas those in *ens*^*swo*^ mutants are under decreased tension compared to controls to confirm that force is applied to nuclei in *bocks*^*DP01391*^ and *klar*^*1*^ mutants but not in *ens*^*swo*^ mutants. More broadly, these data present the first direct evidence that regulation of interactions between nuclei is a critical determinant of nuclear movement and that nucleus-nucleus interactions are LINC complex-dependent. Thus, these data raise the possibility that aligned nuclei in the center of a developing or regenerating muscle are physically linked and that this linkage is critical for nuclear functions.

**Fig. 5.**
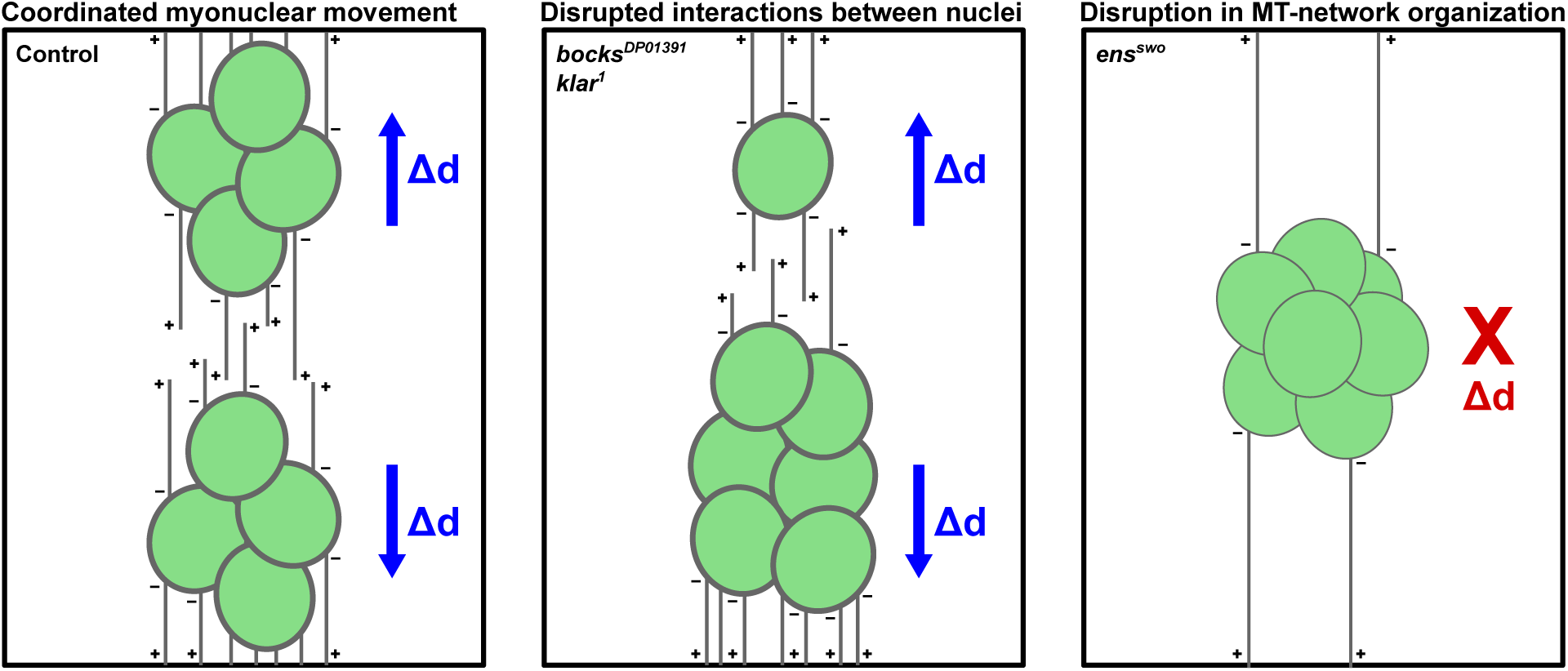
Model of myonuclear movement during *Drosophila* embryonic muscle development. In skeletal muscle, the active translocation of myonuclei (green) is dependent on the integrity of the nuclear envelope and the organization of the microtubule cytoskeleton. To achieve proper nuclear positioning, the two nuclear envelope proteins, bocksbeutel and klarsicht, facilitate the separation and distribution of nuclei into two distinct clusters of equal size by relieving associative interactions between neighboring nuclei. Since each myonucleus acts as a local microtubule organizing center, microtubules (gray) nucleate from the nuclear periphery (minus ends, −) and extend out (plus ends, +) to the cell cortex. These microtubules are able to generate force to pull their attached nuclei via ensconsin, which maintains the organization of the MT-network and promotes the sliding of adjacent microtubules. As a result of the coordinated actions of these proteins, nuclei are pull to the end of the muscle before achieving their final position. Blue arrows denote the direction of net displacement (Δd) of nuclei.

The molecular mechanisms by which klarsicht and bocksbeutel regulate separation of nuclei from their neighbors and the molecular mechanisms by which ensconsin regulates the number of growing microtubules necessitate further investigation. However, we hypothesize that ensconsin may contribute, either directly or indirectly, to microtubule nucleation and anchoring at the nuclear envelope. Recent work found that Bsg25D, the *Drosophila* homolog of Ninein, interacts with ensconsin and that Bsg25D contributed to ensconsin-dependent nuclear positioning (Rosen et al., 2019). Together with our data showing a reduction in the number of growing microtubules, we hypothesize that perhaps Bsg25D is recruiting ensconsin to participate in microtubule nucleation. Alternatively, both Bsg25D and ensconsin may anchor microtubules to the nuclear envelope. Release of microtubule minus ends from the nuclear envelope may potentiate microtubule instability and the reduction in growing microtubules. Indeed, Ninein does contribute to both nucleation and anchoring of microtubules to the centrosome (Delgehyr et al., 2005), and the loss of either function is consistent with the data here and previously published (Rosen et al., 2019).

The molecular mechanism by which bocksbeutel and klarsicht regulate nuclear position is harder to predict. The simplest explanation might be that they are required to recruit microtubule motors as has been seen in other systems (Starr et al., 2001; Wilson and Holzbaur, 2012; Cadot et al., 2012). However, the phenotype seen here is distinct from the phenotypes observed in animals null for either cytoplasmic dynein or kinesin (Folker et al., 2014). Alternatively, work in *C. elegans* found that loss of nucleus anchoring resulted in a similar clustering of nuclei (Starr et al., 2001). But all of the data we present is from developmental stages that require active movement of nuclei rather than anchoring. When combined with our finding that the clusters of nuclei still move in these genotypes, the simplest explanation is that these factors are required for nuclei to separate from one another. Because it is the loss of *bocks* or *klar* that results in the phenotype suggests that either the recruitment of a separation factor or a disruption in cytoskeletal organization is preventing the separation of nuclei. We speculate that this is based on variations in microtubule organization, consistent with our finding that microtubules are asymmetrically organized around nuclei in animals with mutations in either gene. Furthermore, it is likely that the nuclei that emanate from adjacent nuclei can interact with each other and with other nuclei. Thus, the ablation of individual nuclei will ablate the associated microtubule network. Thus, if the molecular glue is either the microtubules directly or indirectly, the data would be similar.

Altogether these data demonstrate that seemingly similar phenotypes are mechanically distinct and provide an approach along with some of the tools necessary to push beyond this basic understanding toward a molecular comprehension of how the movement of many nuclei is coordinated within a single cytoplasm.

## MATERIALS & METHODS

### *Drosophila* genetics

All stocks were grown under standard conditions at 25°C. Stocks used were *apRed* (Richardson et al., 2007), *bocks*^*DP01391*^ (Bloomington Drosophila Stock Center, 21846), *klar*^*1*^ (Bloomington Drosophila Stock Center, 3256), *ens*^*swo*^ (Metzger et al., 2012), and UAS-EB1.eYFP (Rogers et al., 2008). Mutants were balanced and identified using *TM6b, DGY*. The UAS-EB1.eYFP construct was specifically expressed in the mesoderm using the *twist-GAL4, apRed* driver. Flies carrying *apRed* express a nuclear localization signal (NLS) fused to the fluorescent protein DsRed downstream of the *apterous* mesodermal enhancer. This results in the specific labeling of the myonuclei within the lateral transverse (LT) muscles of the *Drosophila* embryo (Richardson et al., 2007). Thus, only nuclei within the LT muscles are labeled using this reporter. The *twist-GAL4, apRed Drosophila* line was made by recombining the *apRed* promoter and the specific *GAL4* driver, with both elements on the second chromosome.

### Immunohistochemistry

Embryos were collected at 25°C and washed in 50% bleach to remove the outer chorion membrane, washed with water, and then fixed in 50% formalin (Sigma, Product # HT501128) diluted in 1:1 heptane for 20 minutes. Embryos were then devitellinized by vortexing in a 1:1 methanol:heptane solution. Primary antibodies for embryo staining were used at the following final dilutions: rabbit anti-DsRed (1:400, Clontech 632496), rat anti-tropomyosin (1:200, Abcam ab50567), mouse anti-GFP (1:50, Developmental Studies Hybridoma Bank GFP-G1). The conjugated fluorescent secondary antibodies used were Alexa Fluor 555 donkey-anti-rabbit (1:200), Alexa Fluor 488 donkey-anti-rat (1:200), and Alexa Fluor 647 donkey-anti-mouse (1:200) (all Life Technologies). Larvae at stage L3 were dissected as previously described (Collins & Mandigo et al., 2017; Auld et al., 2018). In brief, larvae were dissected in ice-cold PIPES dissection buffer containing 100 mM PIPES (Sigma-Aldrich, P6757), 115 mM D-Sucrose (Fisher Scientific, BP220-1), 5 mM Trehalose (Acros Organics, 182550250), 10 mM Sodium Bicarbonate (Fisher Scientific, BP328-500), 75 mM Potassium Chloride (Fisher Scientific, P333-500), 4 mM Magnesium Chloride (Sigma-Aldrich, M1028) and 1 mM EGTA (Fisher Scientific, 28-071-G), then fixed with 10% formalin (Sigma-Aldrich, HT501128). For larval staining, mouse anti-αTubulin (1:200, Sigma-Aldrich T6199) was used. Acti-stain 555 phalloidin (1:400, Cytoskeleton PHDH1-A) and Hoechst 33342 (1 μg/ml) were added with the fluorescent secondary antibody Alexa Fluor 488 donkey-anti-mouse (1:200, Life Technologies). Both embryos and larvae were mounted in ProLong Gold (Life Technologies, P36930).

### Analysis of myonuclear position in *Drosophila* embryos

Embryos at stage 16 were selected to be imaged based on overall embryo shape, the intensity of the apRed and tropomyosin signals, gut morphology, and the morphology of the trachea as previously described (Collins & Mandigo et al., 2017; Auld et al., 2018; Folker et al., 2012). Confocal z-stacks of fixed embryos were acquired on a Zeiss 700 LSM using a Plan-APOCHROMAT 40×, 1.4 NA oil objective with a 1.0× optical zoom. Images were processed as maximum intensity projections and oriented such that top is dorsal, bottom is ventral, left is anterior, and right is posterior. Measurements were made using the Segmented Line tool in Fiji software (Schindelin et al., 2012). Muscle length measurements were taken starting from the dorsal tip and following through the center of each LT muscle, down to the ventral tip. Dorsal and ventral end distances were taken from each LT muscle by measuring the distance between the closest group of nuclei to the dorsal or ventral muscle pole, respectively. Internuclear distances were taken by measuring the shortest distance in between the dorsal and ventral clusters of nuclei within each LT muscle. Internuclear distances were also plotted according to relative frequency. All three measurements are reported as distances normalized to the muscle length (Fig. S1) and as raw values (Fig. S2). All four LT muscles were measured in four hemisegments from each embryo. Statistical analysis was performed with Prism 4.0 (GraphPad).

### Analysis of myonuclear cluster area in *Drosophila* embryos

Area of nuclear clusters were measured in fixed stage 16 embryos as previously described (Collins & Mandigo et al., 2017). In brief, the area of each cluster of nuclei near either the dorsal or ventral muscle pole was measured in Fiji (Schindelin et al., 2012). Total area of nuclear clusters in each LT muscle was calculated by adding the dorsal and ventral areas. The nuclear separation ratio was calculated by dividing the area of the dorsal cluster by the area of the ventral cluster. Nuclear clusters from all four LT muscles were measured in four hemisegments from each embryo. Statistical analysis was performed with Prism 4.0 (GraphPad).

For qualitative nuclear phenotype analysis, embryos were scored on how nuclei were positioned within the first three LT muscles of each hemisegment. LT 4 was excluded for this analysis due to its variable muscle morphology. Nuclear phenotypes were categorized as either “separated; equal distribution” (nuclei properly segregated into two distinct, even clusters with a nuclear separation ratio ≥ 0.85 and ≤ 1.15), “separated; unequal distribution” (nuclei that segregated into two disproportionate clusters with a nuclear separation ratio < 0.85 or > 1.15), “central” (a nucleus that is not associated with either the dorsal or ventral group located in the middle of the myofiber), “clustered” (nuclei remained in a single cluster toward the ventral end of the myofiber), “spread” (nuclei are distributed through the myofiber with no distinct dorsal or ventral clusters) or “swoosh” (nuclei remained in a single cluster within the middle of the myofiber). Linescans of DsRed intensity were performed on 10 LT muscles for each nuclear phenotype and averaged to determine the typical distribution of nuclei in *bocks*^*DP01391*^ and *ens*^*swo*^ genotypes compared to controls.

### Volumetric imaging and analysis of nuclear clusters

Fixed stage 16 embryos were imaged on a Zeiss LSM 880 with Airyscan (super resolution acquisition, 2× Nyquist sampling) using a Plan-APOCHROMAT 40×, 1.3 NA oil objective at a 1.0× optical zoom and 0.15 µm step size interval through the entire depth of the muscle. Post processing of Airyscan images was completed in ZEN Blue 2016 software. Quantitative volumetric analysis was performed in Imaris version 9.2.1 (Bitplane AG). Images were first processed as maximum intensity projections of confocal z-stacks and oriented such that top is dorsal, bottom is ventral, left is anterior, and right is posterior. A volumetric rendering of each nuclear cluster was created using the Surface Visualization tool of the DsRed channel. Volume measurements were automatically computed from the Surface renderings by Imaris. Statistical analysis was performed with Prism 4.0 (GraphPad).

### Live-embryo imaging and analysis

Embryos for live-imaging were prepared as previously described (Collins & Mandigo et al., 2017; Auld et al., 2018). In brief, embryos were collected at 25°C, washed in 50% bleach to remove the outer membrane, washed with water, and mounted with halocarbon oil (Sigma, Product # H8898). For time-lapse imaging of nuclear movement, stage 15 embryos were selected for imaging based on gut morphology, the position of nuclei, and the intensity of the apRed signal as previously described (Collins & Mandigo et al., 2017; Auld et al., 2018; Folker et al., 2012) with the following modifications. Time-lapse images were acquired on a Zeiss 700 LSM using a Plan-APOCHROMAT 40×, 1.4 NA oil objective with a 1.0× optical zoom at an acquisition rate of 1 min/stack for 2 hours. Movies were processed in Fiji (Schindelin et al., 2012) as maximum intensity projections of confocal z-stacks and corrected for drift using the Correct 3D drift plugin. To calculate the separation speed of nuclei, the Line tool was used to measure the distance between dorsal and ventral nuclear clusters at time 0 h and again at time 2 h. Separation speeds were also plotted according to relative frequency. The aspect ratio of ventral clusters was measured at time 0 h using the Shape Descriptors plugin, which calculates aspect ratio of an ellipse by dividing the major axis of the ellipse by its minor axis. An aspect ratio value closer to 1 indicates a more spherical cluster. Tracks following the movement of individual nuclei within clusters were generated using the Manual Tracking plugin. The displacement of each nucleus was calculated as the difference between the final and initial position. Statistical analysis was performed with Prism 4.0 (GraphPad).

To assess for potential fusion defects, the number of nuclei in the LT muscles was counted from live stage 17 embryos when nuclei have separated and maximized their distance from their neighbors. Nuclei within the LT muscles were identified by expression of DsRed. The number of nuclei were counted from all 4 LT muscles within a single hemisegment, with a total of 4 hemisegments analyzed for each embryo.

### 2-photon ablation of myonuclei

Embryos were collected at 25°C and were washed in 50% bleach to remove the outer membrane, washed with water, and mounted with halocarbon oil (Sigma, Product # H8898). Stage 16 embryos were selected for ablation based on gut morphology, the position of nuclei, and the intensity of the apRed signal as previously described (Folker et al., 2012; Collins & Mandigo et al., 2017; Auld et al., 2018). Time-lapse images of embryos before, during, and after ablation were acquired on a Zeiss 710 LSM using a Plan-APOCHROMAT 40×, 1.1 NA water objective with a 1.0× optical zoom at an acquisition rate of 1 s/frame for 30 s. Ablation was performed using the Coherent Chameleon Ultra II femtosecond pulsed-IR laser at 860 nm with 15-17% laser power. As shown in Supplemental Figure 4, a nucleus was selected for ablation by drawing a region of interest (ROI) in ZEN Black 2012 software. For each ablation time-lapse, the first frame (time = 0 s) was taken before the ablation event. The next frame (time = 1 s), shows the ablation of the targeted nucleus, followed by the subsequent response of the remaining nuclei present. Since no muscle marker is present, imaging with transmitted light was used to ensure that ablation did not destroy the surrounding tissue. An ablation was considered successful by the loss of the DsRed signal accompanied by the movement of nuclei. Nuclei that were simply photobleached were characterized by just the loss of DsRed fluorescence without any subsequent response from the embryo (Fig. S4b). A failed ablation attempt that resulted in boiling of the embryo was identified by a hole burned through the membrane (Fig. S4c, arrowhead), as seen through the transmitted light channel.

Movies were processed in Fiji (Schindelin et al., 2012) as single confocal slices and oriented such that top is dorsal, bottom is ventral, left is anterior, and right is posterior. The area of clusters in which a nucleus was ablated was measured before and after the ablation event. The area of nuclear clusters before and after ablation were plotted as a percentage change. The displacement and velocity of nuclear clusters were measured using the centroid measurement, which calculates the center point of a cluster based on the average x and y coordinates of all pixels in the cluster. The total displacement of each cluster was calculated as the cumulative distance traveled over the 30 s after ablation. The initial velocity was defined as the speed a cluster traveled the first second after ablation. Statistical analysis was performed with Prism 4.0 (GraphPad).

### Analysis of microtubule organization in *Drosophila* larvae

Confocal z-stacks of dissected stage L3 larvae were acquired on a Zeiss 700 LSM using a Plan-APOCHROMAT 40×, 1.4 NA oil objective lens at a 0.5× optical zoom for whole muscle images and at a 2.0× optical zoom for regions around myonuclei. Images were processed as maximum intensity projections and oriented such that top is dorsal, bottom is ventral, left is anterior, and right is posterior. Microtubule organization was assessed in two distinct regions of interest within the ventral longitudinal muscle 3 (VL3). The first region consists of microtubules that intersect at regions between nuclei to form a lattice. For these regions, the Texture Detection Technique (TeDT) was used (Liu and Ralston, 2014). TeDT is a robust tool that can assess the orientation of the microtubule network by detecting the dominant angles at which microtubules intersect one another. For TeDT analysis, 200 x 100 square pixel regions of the microtubule lattice that excluded nuclei were cropped from whole muscle images. TeDT analysis on cropped regions was performed in MATLAB (MathWorks) which presented the resulting intersection angles detected as directional histograms (HD) from 0° to 360°.

The second region of interest were microtubules emanating directly from the myonuclei. Polarity of these microtubules was analyzed as previously described (Collins & Mandigo et al., 2017). The fluorescence intensity was measured from a 10 μm x 2 μm region positioned 15 μm anteriorly and 15 μm posteriorly from the center of the nucleus, using the Plot Profile tool in Fiji (Schindelin et al., 2012). Similarly, the fluorescence intensity was also measured from a 2 μm x 10 μm region positioned 15 μm dorsally and 15 μm ventrally from the center of the nucleus. Average fluorescence intensities were calculated for the anterior/posterior (AP) positions as well as the dorsal/ventral (DV) positions. A ratio between the average AP and DV fluorescence intensities was used to determine the microtubule distribution ratio. A value of 1 indicates a uniform distribution of microtubules around the nucleus. Values >1 indicate there are more microtubules distributed within the anterior/posterior regions relative to the nucleus, while values <1 indicate there are more microtubules distributed within the dorsal/ventral regions relative to the nucleus. Organization of microtubules emanating from nuclei was also qualitatively assessed based on the presence of a dense microtubule ring around the nuclear periphery. Images of nuclei were blindly scored for the presence or absence of a microtubule ring. A nucleus was considered to have a microtubule ring based on the contiguous presence of α-tubulin intensity around the perimeter of the nucleus. Statistical analysis was performed with Prism 4.0 (GraphPad).

### Analysis of microtubule dynamics in *Drosophila* embryos

Embryos for live imaging of EB1 comets were collected and prepared similarly. Stage 16 embryos were selected for imaging based on gut morphology, the position of nuclei, and the intensity of the apRed signal as previously described (Collins & Mandigo et al., 2017; Auld et al., 2018; Folker et al., 2012). Time-lapse images of EB1-eYFP were acquired on a Zeiss LSM 880 with Airyscan Fast mode (super resolution acquisition, 2× Nyquist sampling) using a Plan-APOCHROMAT 40×, 1.3 NA oil objective at a 4.0× optical zoom at an acquisition rate of 1 s/frame for 60 s. Post processing of Airyscan Fast images was done in ZEN Blue 2016 software. EB1 comets were imaged within the LT muscles as well as the dorsal oblique (DO) muscles, which are a flatter muscle group, ideal for imaging quick dynamics. Movies were processed as single confocal slices in Fiji (Schindelin et al., 2012). Time-lapse images taken in the LT muscles were oriented such that top is dorsal, bottom is ventral, left is anterior, and right is posterior. Time-lapse images taken in the DO muscles were oriented such that top is posterior, bottom is anterior, left is dorsal, and right is ventral. Trajectories of EB1 comets were made from time-lapse images using the Temporal-Color Code plugin, which sums up the first 15 consecutive frames (1 s each), and then overlays the resulting image to a blue-green-red color sequence, with each color representing a total of 5 seconds. All quantifications of EB1 dynamics was performed on temporal overlays by hand. Only comets that were visible for the full 15 seconds were used in this analysis. The starting position of each comet was categorized within the LT muscles as either starting within the dorsal pole region, ventral pole region, or between nuclei. Similarly, the starting position of each comet was categorized within the DO muscles as either starting within the anterior pole region, posterior pole region, or between nuclei. The direction of EB1 comets was also determined as either traveling dorsally/posteriorly or ventrally/anteriorly and whether the comets move toward or away from the nearest myotendinous junction. The length of EB1 trajectories over the 15 s timeframe was measured to calculate EB1 comet velocity over the 1 min time-lapse. The number of EB1 comets was counted and normalized to the muscle area. Statistical analysis was performed with Prism 4.0 (GraphPad).

## Acknowledgments

We thank Wendy C. Salmon and the W.M. Keck Microscopy Facility at the Whitehead Institute for infrastructure and support with regard to all 2-photon laser ablation experiments. Additionally, we thank Bret Judson and the Boston College Imaging Core for infrastructure and support with regard to super-resolution imaging using the Zeiss LSM 880 Airyscan microscope. *Drosophila* stocks obtained from the Bloomington Drosophila Stock Center (NIHP400D018537) were used in this study.

## Author contributions

Conceptualization: M.A.C., E.S.F.; Methodology: M.A.C., E.S.F.; Formal analysis: M.A.C., L.A.C., R.T.; Investigation: M.A.C., L.A.C., R.T., T.R.M., E.W.; Resources: E.S.F.; Writing - original draft: M.A.C.; Writing - review & editing: M.A.C., E.S.F.; Visualization: M.A.C.; Supervision: E.S.F.; Project administration: E.S.F.; Funding acquisition: E.S.F.

## Data and material availability

All *Drosophila* stocks are available upon request. All data necessary for confirming the conclusions of the article are present within the article, figures, and tables.

## Supplementary Information

**Fig. S1.**
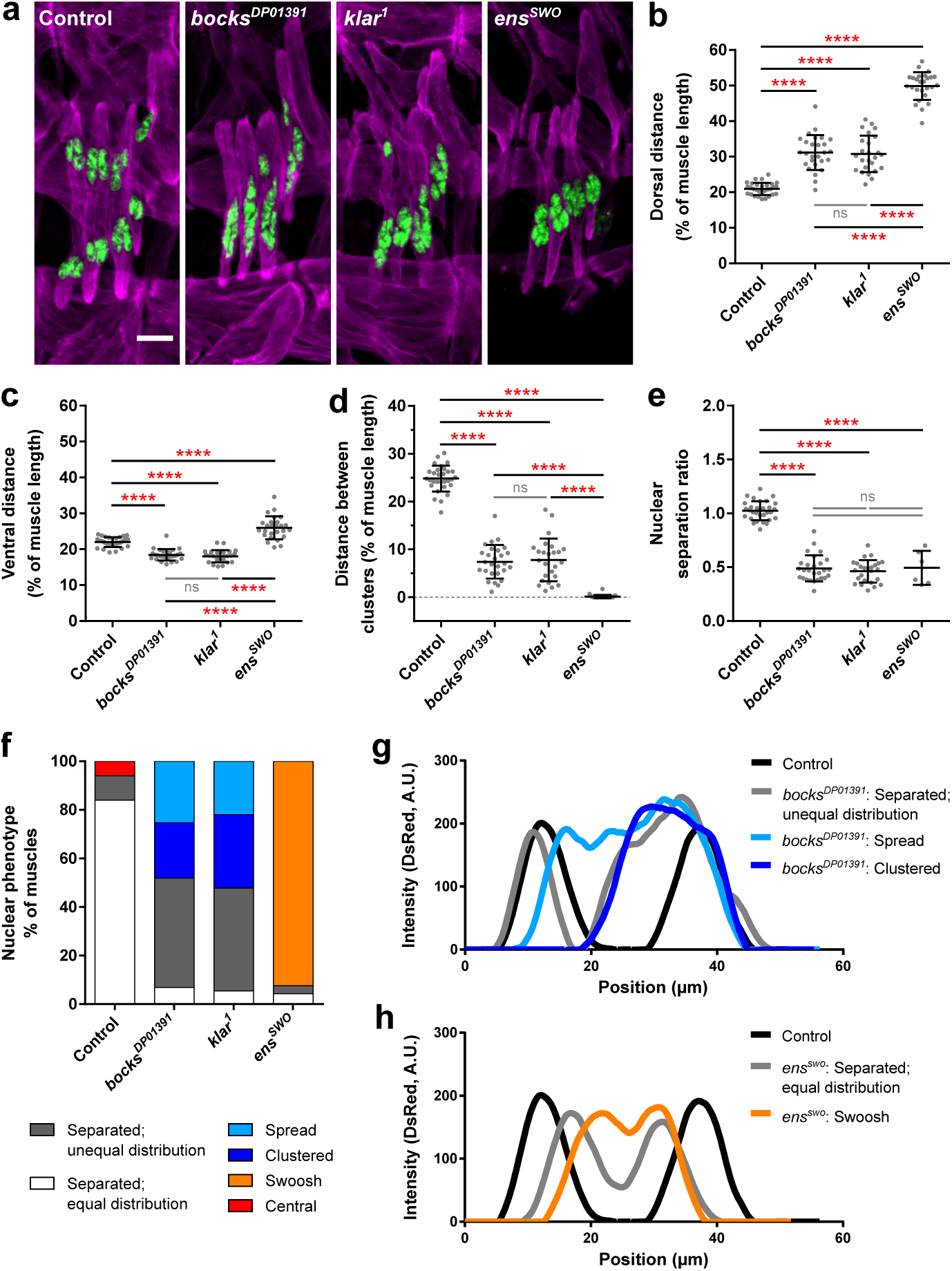
Bocksbeutel, klarsicht, and ensconsin regulate myonuclear position in *Drosophila* embryos. (**a**) Immunofluorescence images of the lateral transverse muscles in one hemisegment from stage 16 (16 hours AEL) embryos for the indicated genotypes. Muscles in magenta, myonuclei in green. Scale bar, 10 µm. (**b–d**) Graphs indicating the distance between the dorsal end of the muscle and the nearest nucleus (**b**), the distance between the ventral end of the muscle and the nearest nucleus (**c**), and the distance between the dorsal and ventral clusters of nuclei (**d**). All distances were normalized to the muscle length. (**e**) The relative size of the dorsal cluster of nuclei compared to the ventral cluster of nuclei. It is important to note that in 21 out of the 27 *ens*^*swo*^ embryos, there was only one cluster present. Thus, the nuclear separation ratio was only calculated for the 6 embryos that had two distinct clusters. Data points in (b–e) correspond to the average value within a single embryo. Error bars indicate the s.d. from ≥25 embryos for each genotype taken from at least three independent experiments. One-way ANOVA with Tukey HSD post hoc test was used to assess the statistical significance of differences in measurements between all experimental groups. (**f**) The frequency at which each nuclear positioning phenotype was observed in each of the indicated genotypes. (**g–h**) Averaged linescans of DsRed intensity for each nuclear phenotype observed in *bocks*^*DP01391*^ mutants (**g**) and *ens*^*swo*^ mutants (**h**) compared to controls. Position correlates to the length of the muscle. Dorsal end position corresponds to 0 μm.

**Fig. S2.**
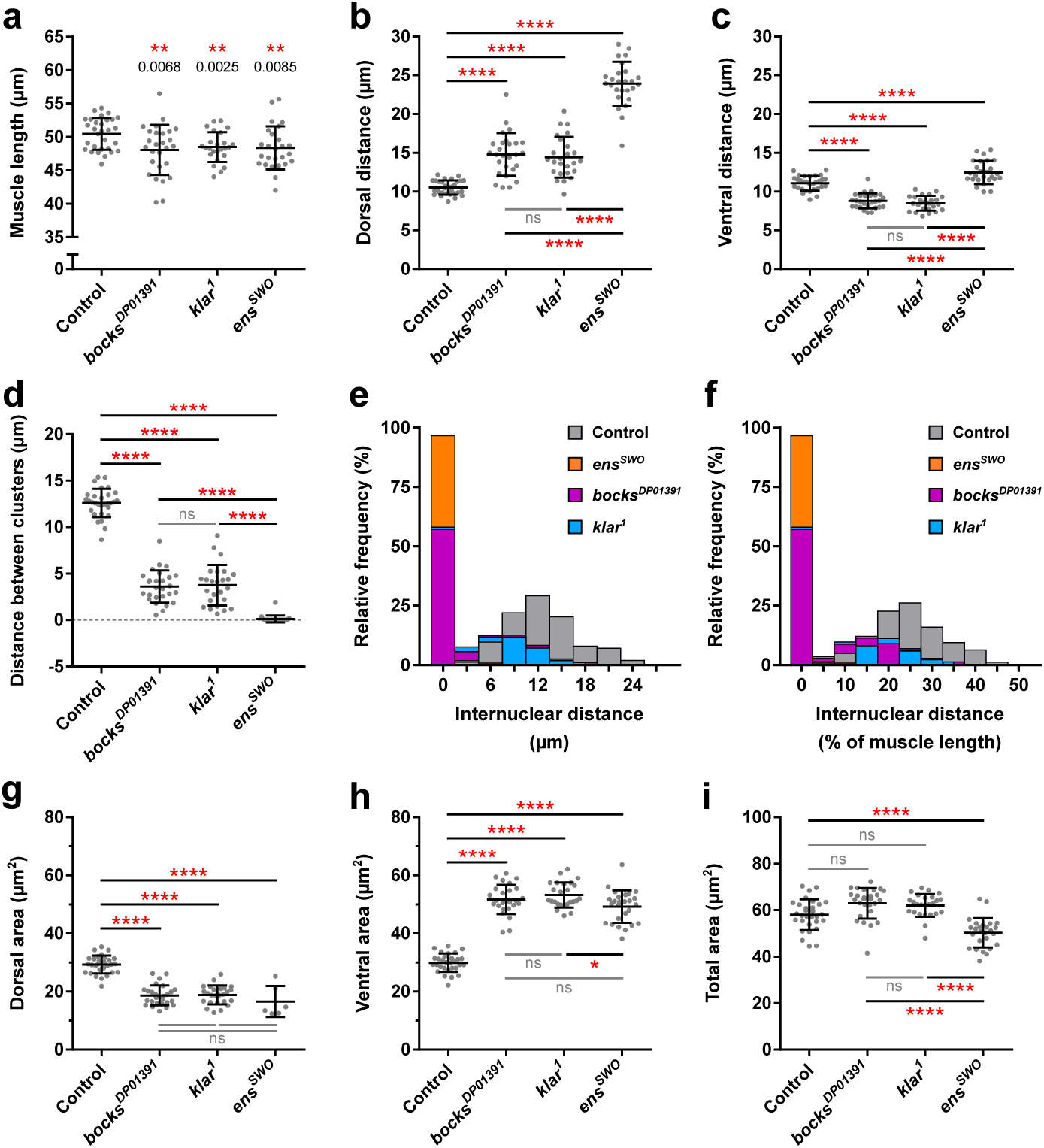
Bocksbeutel, klarsicht, and ensconsin are necessary for proper muscle length and myonuclear position in *Drosophila* embryos. (**a**) The average length of the lateral transverse muscles for the indicated genotypes. (**b–d**) Graphs indicating the raw distance between the dorsal end of the muscle and the nearest nucleus (**b**), the raw distance between the ventral end of the muscle and the nearest nucleus (**c**), and the raw distance between the dorsal and ventral nuclear clusters (**d**). (**e–f**) The relative distribution of all internuclear distances measured, represented as raw values (**e**) and as a function of muscle length (**f**). (**g–i**) Graphs indicating the area of nuclei located near the dorsal end of the muscle (**g**), the area of nuclei located near the ventral end of the muscle (**h**), and the total area of all myonuclei present within the muscle (**i**). It is important to note that in 21 out of the 27 *ens*^*swo*^ embryos, there was only one cluster present. Thus, the dorsal cluster area was only measured in the 6 embryos that had two distinct clusters. Data points in (a–d) and (g–i) correspond to the average value within a single embryo. Error bars indicate the s.d. from ≥25 embryos for each genotype taken from at least three independent experiments. For (a) Student’s t-test with Welsh’s correction was used to assess the statistical significance of differences in measurements between experimental genotypes to controls. For (b–d) and (g–i) One-way ANOVA with Tukey HSD post hoc test was used to assess the statistical significance of differences in measurements between all experimental groups.

**Fig. S3.**
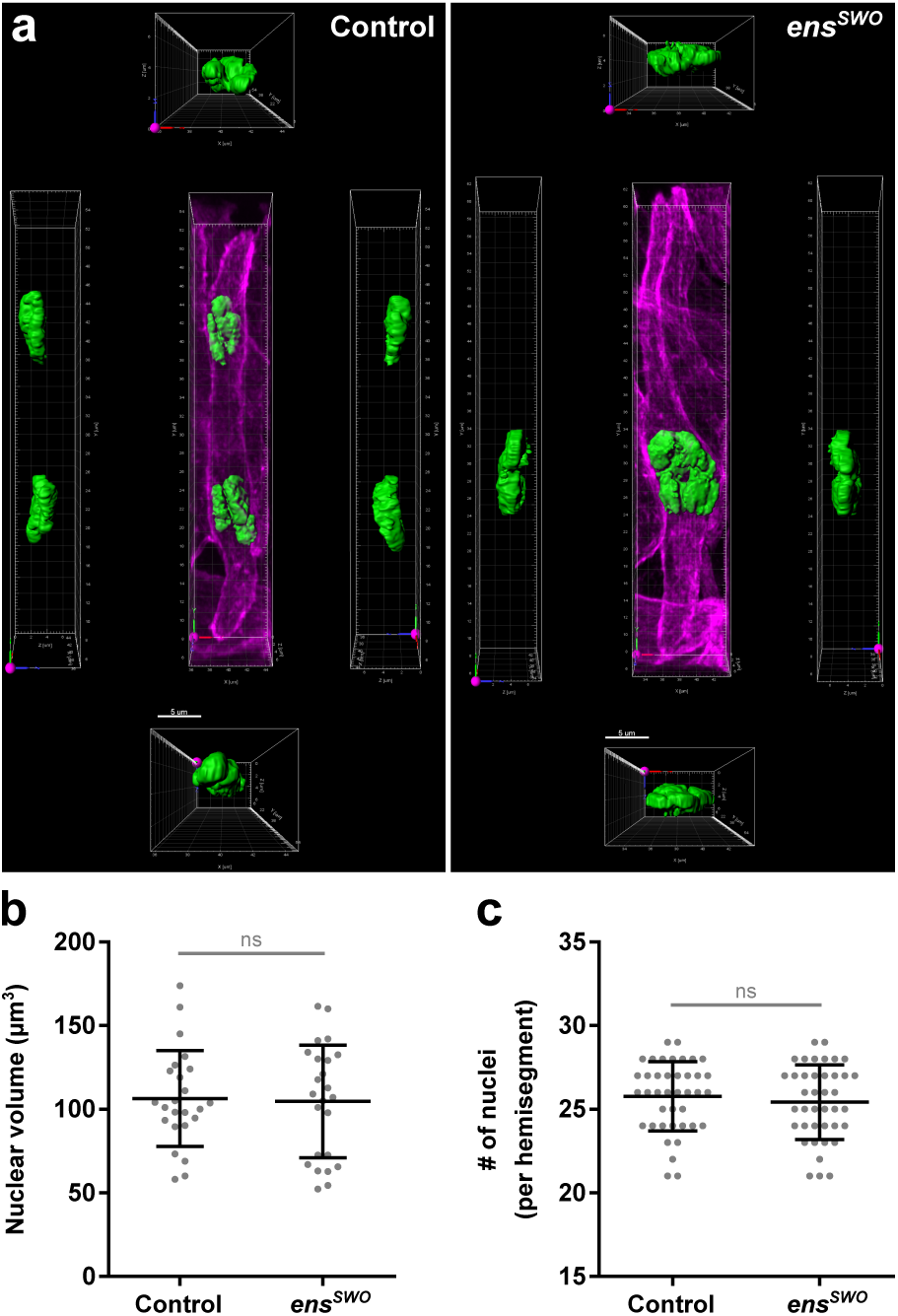
Total nuclear volume and number of nuclei are not disrupted in *ensconsin*-depleted embryos. (**a**) Three-dimensional volumetric renderings of nuclear clusters created from Airyscan images of a single LT muscle from stage 16 (16 hours AEL) control and *ens*^*swo*^ embryos. Muscles in magenta, myonuclei in green. Scale bar, 5 µm. Each rendering showing just the nuclei have been rotated -90° (left) and +90° (right) along the y-axis as well as -90° (bottom) and +90° (top) along the x-axis, relative to the center image. (**b**) The total volume of nuclei within a single LT muscle. Data points correspond to the total volume of nuclei within a single LT muscle. Error bars indicate the s.d. from 24 LT muscles for each genotype measured from six different embryos. Student’s t-test with Welsh’s correction was used to assess the statistical significance of differences in nuclear volume between *ens*^*swo*^ embryos and controls. (**c**) The number of nuclei per hemisegment counted from live stage 17 (17 hours AEL) control and *ens*^*swo*^ embryos. Data points correspond to the total number of nuclei counted within a single hemisegment. Error bars indicate the s.d. from 40 hemisegments for each genotype taken from 10 different embryos. Student’s t-test with Welsh’s correction was used to assess the statistical significance of differences in the number of nuclei counted from *ens*^*swo*^ embryos and controls.

**Fig. S4.**
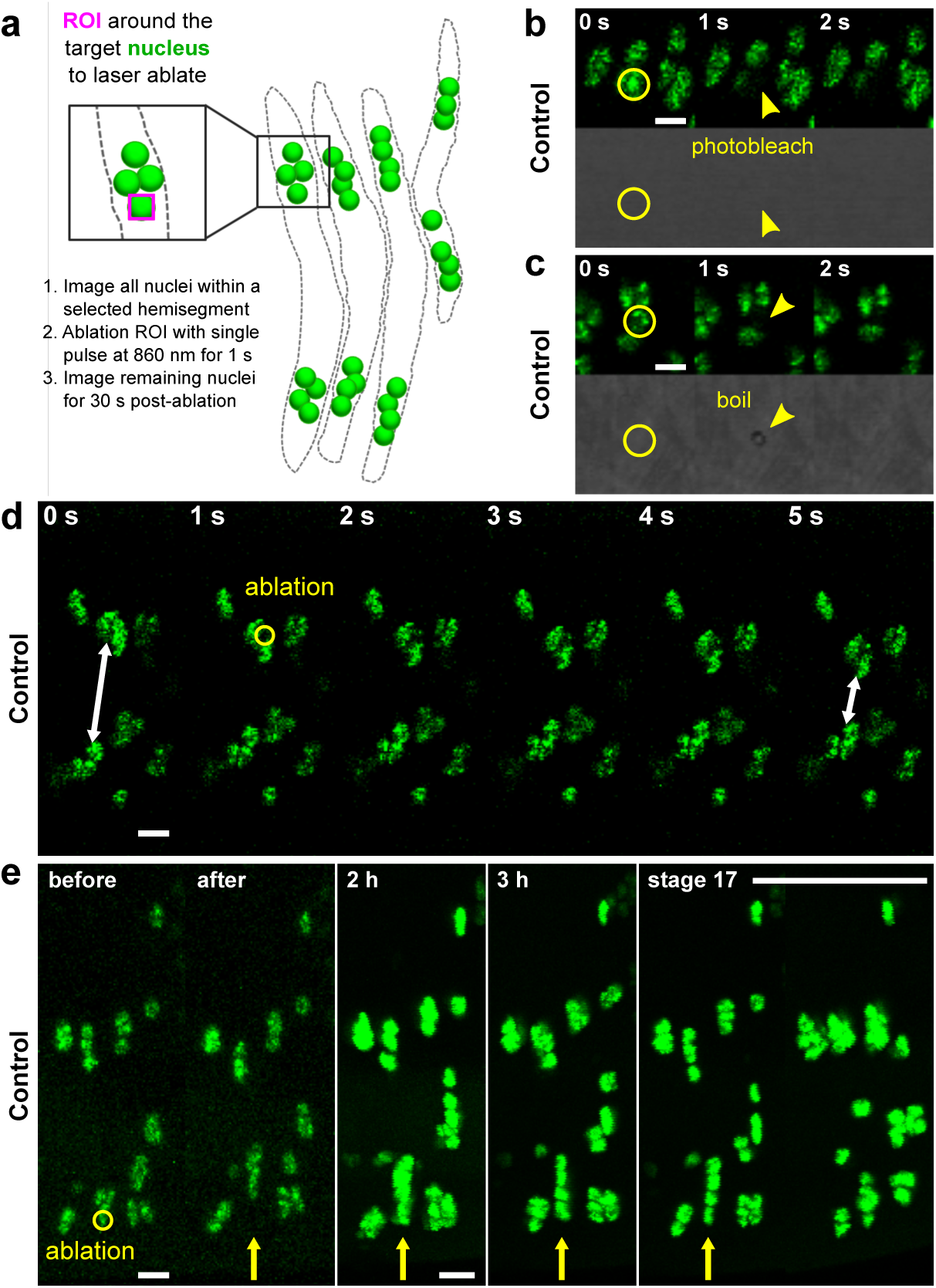
*In vivo* 2-photon laser ablation of myonuclei within *Drosophila* embryos. (**a**) Schematic illustrating how myonuclei are ablated in the lateral transverse (LT) muscles of a living stage 16 (16 hours AEL) control embryo. Nuclei (green) in the LT muscles (dotted grey outline) are identified by the expression of DsRed. Before ablation, all nuclei within a hemisegment are imaged. The nucleus to be ablated is selected by a region of interest (magenta ROI) and then ablated using a pulsed 2-photon laser at 860 nm for 1 s. The remaining nuclei are then imaged every second for 30 s to observe the post-ablation response. (**b–c**) Montages from time-lapse images showing failed ablation attempts. Nuclei in green, transmitted light in gray. Photobleached nuclei were characterized by just the loss of fluorescence with no subsequent response (**b**) while embryos that were boiled were identified by a hole burned through the membrane (**c**, arrowhead). Scale bar, 5 µm. (**d**) Montage from a time-lapse image showing the ablation of a single nucleus within the LT muscles of a stage 16 control embryo. The first frame shows all the nuclei before the ablation event (0 s). The next frame (1 s) shows the ablation of a single nucleus (yellow circle), followed by the subsequent response of the remaining nuclei present within the cluster after the ablation event (white arrows). (**e**) Still images from a stage 16 embryo that was followed from the time of ablation until stage 17 (the final embryonic stage) to demonstrate that ablation does not affect embryonic development or viability. Scale bar, 10 µm.

**Table S1.** Summary of P-values. The following scale was used to determine statistical significance: not significant (ns) ≥ 0.05, *P<0.05, **P<0.01, ***P<0.001, ****P<0.0001, N/A not applicable.

| Statistical Analysis | Genotype Comparisons |  |  |  |  |  |
| --- | --- | --- | --- | --- | --- | --- |
|  | control<br>vs.<br>*bocks*<sup>DP01391</sup> | control<br>vs.<br>*klar*<sup>1</sup> | control<br>vs.<br>*ens*<sup>SWO</sup> | *bocks*<sup>DP01391</sup><br>vs.<br>*klar*<sup>1</sup> | *bocks*<sup>DP01391</sup><br>vs.<br>*ens*<sup>SWO</sup> | *klar*<sup>1</sup><br>vs.<br>*ens*<sup>SWO</sup> |
| **Figure 1** (One-way ANOVA with Tukey HSD post hoc test) |  |  |  |  |  |  |
| **1b:** Separation speed | **0.0003** | **0.0029** | **< 0.0001** | 0.9957 | **< 0.0001** | **< 0.0001** |
| **1d:** Ventral cluster aspect ratio | **< 0.0001** | **< 0.0001** | 0.9728 | 0.9200 | **< 0.0001** | **< 0.0001** |
| **1f:** Displacement | 0.3952 | 0.7351 | **< 0.0001** | 0.0655 | **< 0.0001** | **< 0.0001** |
| **Figure 2** (One-way ANOVA with Tukey HSD post hoc test) |  |  |  |  |  |  |
| **2b:** Nuclear cluster area (before) | **0.0002** | **0.0002** | 0.1347 | >0.9999 | **0.0155** | **0.0134** |
| **2b:** Nuclear cluster area (after) | 0.0861 | 0.0851 | **0.0050** | >0.9999 | 0.4740 | 0.4783 |
| **2b':** % change in cluster area | **<0.0001** | **<0.0001** | **0.0081** | 0.9989 | **<0.0001** | **<0.0001** |
| **2c':** Total cluster displacement | **0.0004** | **0.0006** | **0.0044** | 0.9954 | **<0.0001** | **<0.0001** |
| **2d':** Initial velocity (V<sub>0</sub>) | **0.0039** | **0.0342** | **0.0003** | 0.7175 | **<0.0001** | **<0.0001** |
| **Figure 3** (One-way ANOVA with Tukey HSD post hoc test) |  |  |  |  |  |  |
| **3e:** MT Distribution (AP:DV) | **< 0.0001** | **< 0.0001** | **< 0.0001** | 0.8361 | **<0.0001** | **0.0002** |
| **Figure 4** (Student's t-test with Welch's correction) |  |  |  |  |  |  |
| **4d:** EB1 comet velocity (LT muscles) | N/A | N/A | 0.4296 | N/A | N/A | N/A |
| **4d:** EB1 comet velocity (DO muscles) | N/A | N/A | 0.5967 | N/A | N/A | N/A |
| **4e:** # of EB1 comets (LT muscles) | N/A | N/A | **0.0024** | N/A | N/A | N/A |
| **4e:** # of EB1 comets (DO muscles) | N/A | N/A | **0.0094** | N/A | N/A | N/A |
| **Supplemental Figure 1** (One-way ANOVA with Tukey HSD post hoc test) |  |  |  |  |  |  |
| **S1b:** Dorsal distance (%) | **< 0.0001** | **< 0.0001** | **< 0.0001** | 0.9858 | **< 0.0001** | **< 0.0001** |
| **S1c:** Ventral distance (%) | **< 0.0001** | **< 0.0001** | **< 0.0001** | 0.8985 | **< 0.0001** | **< 0.0001** |
| **S1d:** Internuclear distance (%) | **< 0.0001** | **< 0.0001** | **< 0.0001** | 0.9685 | **< 0.0001** | **< 0.0001** |
| **S1e:** Nuclear separation ratio | **< 0.0001** | **< 0.0001** | **< 0.0001** | 0.8120 | 0.9995 | 0.9142 |
| **Supplemental Figure 2** (*Student's t-test with Welch's correction; One-way ANOVA with Tukey HSD post hoc test) |  |  |  |  |  |  |
| **S2a:** Muscle length* | **0.0068** | **0.0024** | **0.0085** | N/A | N/A | N/A |
| **S2b:** Dorsal distance (μm) | **< 0.0001** | **< 0.0001** | **< 0.0001** | 0.9462 | **< 0.0001** | **< 0.0001** |
| **S2c:** Ventral distance (μm) | **< 0.0001** | **< 0.0001** | **< 0.0001** | 0.7650 | **< 0.0001** | **< 0.0001** |
| **S2d:** Internuclear distance (μm) | **< 0.0001** | **< 0.0001** | **< 0.0001** | 0.9878 | **< 0.0001** | **< 0.0001** |
| **S2g:** Dorsal area | **< 0.0001** | **< 0.0001** | **< 0.0001** | 0.9980 | 0.5315 | 0.4700 |
| **S2h:** Ventral area | **< 0.0001** | **< 0.0001** | **< 0.0001** | 0.6108 | 0.2171 | **0.0123** |
| **S2i:** Total area | 0.0519 | 0.0888 | **< 0.0001** | 0.9477 | **< 0.0001** | **< 0.0001** |
| **Supplemental Figure 3** (Student's t-test with Welch's correction) |  |  |  |  |  |  |
| **S3b:** Nuclear volume | N/A | N/A | 0.8487 | N/A | N/A | N/A |
| **S3c:** # of nuclei | N/A | N/A | 0.4689 | N/A | N/A | N/A |

**Movie S1. Volumetric imaging of myonuclei in the lateral transverse muscle of a control *Drosophila* embryo**.

Movie of a three-dimensional volumetric rendering of the dorsal and ventral nuclear clusters within a single LT muscle from a stage 16 (16 hours AEL) control embryo. Muscles in magenta, myonuclei in green. Scale bar, 5 µm. The LT muscle is rotated 360° along the x-axis and 360° along the y-axis.

**Movie S2. Volumetric imaging of myonuclei in the lateral transverse muscle of an *ens***^***swo***^ **mutant embryo**.

Movie of a three-dimensional volumetric rendering of the nuclear cluster within a single LT muscle from a stage 16 (16 hours AEL) *ens*^*swo*^ embryo. Muscles in magenta, myonuclei in green. Scale bar, 5 µm. The LT muscle is rotated 360° along the x-axis and 360° along the y-axis.

**Movie S3. Nuclear migration in the lateral transverse muscle of a control *Drosophila* embryo**.

Time-lapse acquisition showing the migration of myonuclei within four lateral transverse (LT) muscles of a control embryo. Tracks correspond to the movement of individual nuclei within each cluster over the course of two hours. Time-lapse starts at stage 15 (15 hours AEL, t = 0 min), when nuclei have already separated into two distinct clusters. Each LT muscle has one dorsal cluster and one ventral cluster that migrate directionally to opposite ends of the muscle. At stage 16 (16 hours AEL), the dorsal and ventral clusters have reached their respective muscle pole, maximizing the distance between them. Scale bar, 10 µm.

**Movie S4. Altered nuclear migration in the lateral transverse muscle of a *bocks***^***DP01391***^ **mutant embryo**.

Time-lapse acquisition showing the migration of myonuclei within four lateral transverse (LT) muscles of a *bocks*^*DP01391*^ mutant embryo. Tracks correspond to the movement of individual nuclei over the course of two hours. Time-lapse starts at stage 15 (15 hours AEL, t = 0 min), where a majority of nuclei failed to separate and remain clustered together in the ventral end of the muscle. Only two escaper nuclei separate from the ventral cluster and migrate directionally toward the dorsal muscle pole. Scale bar, 10 µm.

**Movie S5. Altered nuclear migration in the lateral transverse muscle of a *klar***^***1***^ **mutant embryo**.

Time-lapse acquisition showing the migration of myonuclei within four lateral transverse (LT) muscles of a *klar*^*1*^ mutant embryo. Tracks correspond to the movement of individual nuclei over the course of two hours. Time-lapse starts at stage 15 (15 hours AEL, t = 0 min), where a majority of nuclei failed to separate and remain clustered together in the ventral end of the muscle. Only one escaper nucleus separates from the ventral cluster and migrates directionally toward the dorsal muscle pole. Scale bar, 10 µm.

**Movie S6. Altered nuclear migration in the lateral transverse muscle of an *ens***^***swo***^ **mutant embryo**.

Time-lapse acquisition showing the migration of myonuclei within four lateral transverse (LT) muscles of an *ens*^*swo*^ mutant embryo. Tracks correspond to the movement of individual nuclei over the course of two hours. Time-lapse starts at stage 15 (15 hours AEL, t = 0 min). In each LT muscle, none of the nuclei separate and remain within a single cluster. Scale bar, 10 µm.

**Movie S7. *In vivo* 2-photon laser ablation of myonuclei in a control *Drosophila* embryo**.

Time-lapse acquisition showing the ablation of a myonucleus within the lateral transverse (LT) muscles of a stage 16 (16 hours AEL) control embryo. The first frame shows the nuclei before ablation (0 s). The next frame (1 s) shows the ablation of a single nucleus (yellow circle), followed by the subsequent response of the remaining nuclei after ablation (2-5 s). Myonuclei in green, transmitted light in gray. Scale bar, 10 µm.

**Movie S8. *In vivo* 2-photon laser ablation of myonuclei in a *bocks***^***DP01391***^ **mutant embryo**.

Time-lapse acquisition showing the ablation of a myonucleus within the lateral transverse (LT) muscles of a stage 16 (16 hours AEL) *bocks*^*DP01391*^ mutant embryo. The first frame shows the nuclei before ablation (0 s). The next frame (1 s) shows the ablation of a single nucleus (yellow circle), followed by the subsequent response of the remaining nuclei after ablation (5-30 s). Myonuclei in green, transmitted light in gray. Scale bar, 10 µm.

**Movie S9. *In vivo* 2-photon laser ablation of myonuclei in a *klar***^***1***^ **mutant embryo**.

Time-lapse acquisition showing the ablation of a myonucleus within the lateral transverse (LT) muscles of a stage 16 (16 hours AEL) *klar*^*1*^ mutant embryo. The first frame shows the nuclei before ablation (0 s). The next frame (1 s) shows the ablation of a single nucleus (yellow circle), followed by the subsequent response of the remaining nuclei after ablation (5-30 s). Myonuclei in green, transmitted light in gray. Scale bar, 10 µm.

**Movie S10. *In vivo* 2-photon laser ablation of myonuclei in an *ens***^***swo***^ **mutant embryo**.

Time-lapse acquisition showing the ablation of a myonucleus within the lateral transverse (LT) muscles of a stage 16 (16 hours AEL) *ens*^*swo*^ mutant embryo. The first frame shows the nuclei before ablation (0 s). The next frame (1 s) shows the ablation of a single nucleus (yellow circle), followed by the subsequent response of the remaining nuclei after ablation (5-30 s). Myonuclei in green, transmitted light in gray. Scale bar, 10 µm.

**Movie S11. *In vivo* imaging of EB1 comet dynamics in the lateral transverse muscles of a control *Drosophila* embryo**.

Time-lapse acquisition of the lateral transverse muscles in a stage 16 (16 hours AEL) control embryo expressing EB1.eYFP. Time course, 60 s. Scale bar, 5 μm.

**Movie S12. *In vivo* imaging of EB1 comet dynamics in the lateral transverse muscles of an *ens***^***swo***^ **mutant embryo**.

Time-lapse acquisition of the lateral transverse muscles in a stage 16 (16 hours AEL) *ens*^*swo*^ mutant embryo expressing EB1.eYFP. Time course, 60 s. Scale bar, 5 μm.

**Movie S13. *In vivo* imaging of EB1 comet dynamics in the dorsal oblique muscles of a control *Drosophila* embryo**.

Time-lapse acquisition of the dorsal oblique muscles in a stage 16 (16 hours AEL) control embryo expressing EB1.eYFP. Time course, 60 s. Scale bar, 5 μm.

**Movie S14. *In vivo* imaging of EB1 comet dynamics in the dorsal oblique muscles of an *ens***^***swo***^ **mutant embryo**.

Time-lapse acquisition of the dorsal oblique muscles in a stage 16 (16 hours AEL) *ens*^*swo*^ mutant embryo expressing EB1.eYFP. Time course, 60 s. Scale bar, 5 μm.

